# De novo synthesis and salvage pathway coordinately regulates polyamine homeostasis and determines T cell proliferation and function

**DOI:** 10.1101/2020.04.29.068759

**Authors:** Ruohan Wu, Xuyong Chen, Siwen Kang, Tingting Wang, JN Rashida Gnanaprakasam, Yufeng Yao, Lingling Liu, Song Guo Zheng, Gaofeng Fan, Mark R Burns, Ruoning Wang

**Author notes:** Wu R, Chen X and Kang S contributed equally to this paper. Correspondence should be addressed to: Ruoning Wang.

## Abstract

Robust and effective T cell-mediated immune responses require proper allocation of metabolic resources through metabolic pathways to sustain the energetically costly immune response. As an essential class of polycationic metabolites ubiquitously present in all living organisms, the polyamine pool is tightly regulated by biosynthesis and salvage pathway. We demonstrated that arginine is a major carbon donor and glutamine is a minor carbon donor for polyamine biosynthesis in T cells. Accordingly, the dependence of T cells can be partially relieved by replenishing the polyamine pool. In response to the blockage of de novo synthesis, T cells can rapidly restore the polyamine pool through a compensatory increase in polyamine uptake from the environment, indicating a layer of metabolic plasticity. Simultaneously blocking synthesis and uptake depletes the intracellular PA pool, inhibits T cell proliferation, suppresses T cell inflammation, indicating the potential therapeutic value of targeting the polyamine for managing inflammatory and autoimmune diseases.

## Introduction

To cope with pathogens’ capacity for exponential growth, T cells have evolved mechanisms for rapidly adjusting their metabolism in response to T cell receptor (TCR) activation and additional signals indicating environmental change. A robust and effective metabolic reprogramming enables T cells to rapidly expand in number and differentiate to achieve large numbers of effector cells specialized in producing high levels of cytokines. Emerging evidences have shown that metabolic rewiring of the central carbon metabolism maximizes the acquisition and assimilation of energy and carbon, and prepare T cells for growth, differentiation, immune regulation and defense (*1*–*7*). Glycolysis, the pentose phosphate pathway (PPP), the Krebs cycle and fatty acid oxidation (FAO) represent a core set of metabolic pathways that transform carbon and chemical energy from environmental nutrients to support the bioenergetic and biosynthesis needs of T cells. Beyond that, a myriad of peripheral metabolic pathways is integrated to complex metabolic networks and are tightly regulated to generate specialized metabolites, which are essential for maintaining homeostasis and immune functions of T cells.

Non-essential amino acids including arginine, glutamine, and proline have both anabolic and catabolic functions, providing building blocks for protein and connecting central carbon metabolism to a variety of specialized metabolism, including polyamine biosynthesis (*8*–*13*). Polyamine is an essential class of polycationic metabolites ubiquitously present in all living organisms. In addition to de novo biosynthesis, other metabolic routes including polyamine catabolism, influx and efflux act in concert to determine the size of the intracellular polyamine pool (*14*, *15*). Disrupting polyamine homeostasis can affect a plethora of cellular processes, including transcription, translation, redox balance, and mitochondria quality control (*16*–*18*). Dysregulation of the level of polyamine and its amino acid precursors has been found to be associated with inflammation and autoimmune diseases (*18*–*23*). We have previously reported that polyamine is one of the most upregulated metabolite groups following T cell activation, and transcription factor MYC is responsible for its upregulation (*24*). Emerging evidence have also shown that polyamine homeostasis is tightly regulated in cellular contexts other than T cells, which have critical roles in immune regulation and defense (*23*–*30*). As such, a better understanding of how polyamine homeostasis is regulated in immune cells will reveal the fundamental principles of the emerging connections between immune cell metabolic fitness and functional robustness. Further knowledge will also enable us to devise rational and practical approaches to treat inflammatory and autoimmune diseases.

Intrinsic T cell signaling cascades are instrumental in the control of T cell metabolic programming (*31*–*36*). Numerous extrinsic environmental factors including oxygen and nutrient supplies also significantly influence T cell metabolic phenotypes and, thus, immune functions *in vivo* (*3*, *37*). Here, we report that the intracellular polyamine pool is tightly regulated by de novo biosynthesis and salvage, through the import of extracellular polyamine. Heightened arginine and glutamine catabolism provide carbon sources to support polyamine de novo biosynthesis *in vitro*. Genetic and pharmacologic ablation of de novo biosynthesis of polyamine is sufficient to deplete the polyamine pool and suppress T cell proliferation *in vitro*. However, de novo biosynthesis is dispensable in driving T cell proliferation *in vivo* where T cells can salvage circulating polyamine to maintain intracellular polyamine pool. Simultaneously blocking polyamine synthesis and salvage inhibits T cell proliferation *in vivo* and confers protection against the pathogenic development of experimental autoimmune encephalomyelitis (EAE). Our findings implicate the potential therapeutic value of targeting the polyamine metabolism in treating and managing inflammatory and autoimmune diseases.

## Results

### Inhibition of ODC reduces T cell proliferation and viability *in vitro*

We previously reported that a Myc-dependent non-canonical metabolic pathway links amino acid catabolism to the biosynthesis of polyamine during T cell activation (*38*). To investigate the role of polyamine metabolism in T cells, we employed a genetic and a pharmacologic approach to ablate polyamine de novo biosynthesis. Since ornithine decarboxylase (ODC), the rate-limiting enzyme in polyamine de novo biosynthetic pathway, is essential for early embryo development, and ODC germline knockout is embryonically lethal (*39*), we obtained a mouse strain containing a reporter-tagged conditional allele of ODC (FRT-LacZ; ODC^fl^) generated by the European Mouse Mutant Archive (*40*). We first crossed this strain with the FLP knock-in mouse strain, which removed the LacZ-reporter allele and generated the strain containing the conditional allele (ODC^fl^). Then, we generated a T cell-specific ODC knockout strain (ODC cKO) by crossing the ODC^fl^ strain with the CD4-Cre strain. The deletion of ODC was validated by qPCR (Fig. S1A), and the ablation of polyamine de novo synthesis was further validated by the accumulation of ornithine (substrate of ODC) and the depletion of spermine, spermidine, putrescine and N-Acetylputrescine (Fig. S1B). ODC deletion did not affect the distribution of T cell subsets in the thymus, spleen and lymph nodes (Fig. S2A and S2B). In addition, the percentage of naturally occurring IFN-γ-producing, IL-17-producing, and FoxP3^+^ CD4 T cells is comparable in both WT and ODC cKO animals (Fig. S2C). However, genetic deletion of ODC significantly delayed cell cycle progression from G0/G1 to the S phase after T cell activation and suppressed overall T cell proliferation in vitro (Fig. 1A and 1B). Consistent with the impact of genetic deletion of ODC on T cells, difluoromethylornithine (DFMO), a potent inhibitor of ODC, inhibited activation-induced T cell cycle progression and proliferation in vitro (Fig. 1C and 1D). Finally, both genetic deletion of ODC and DFMO treatment caused moderately more cell death after activation in a time-dependent manner (Fig. 1E and 1F). Together, our results indicate that polyamine homeostasis is critical for T cell proliferation and survival.

**Figure 1.**
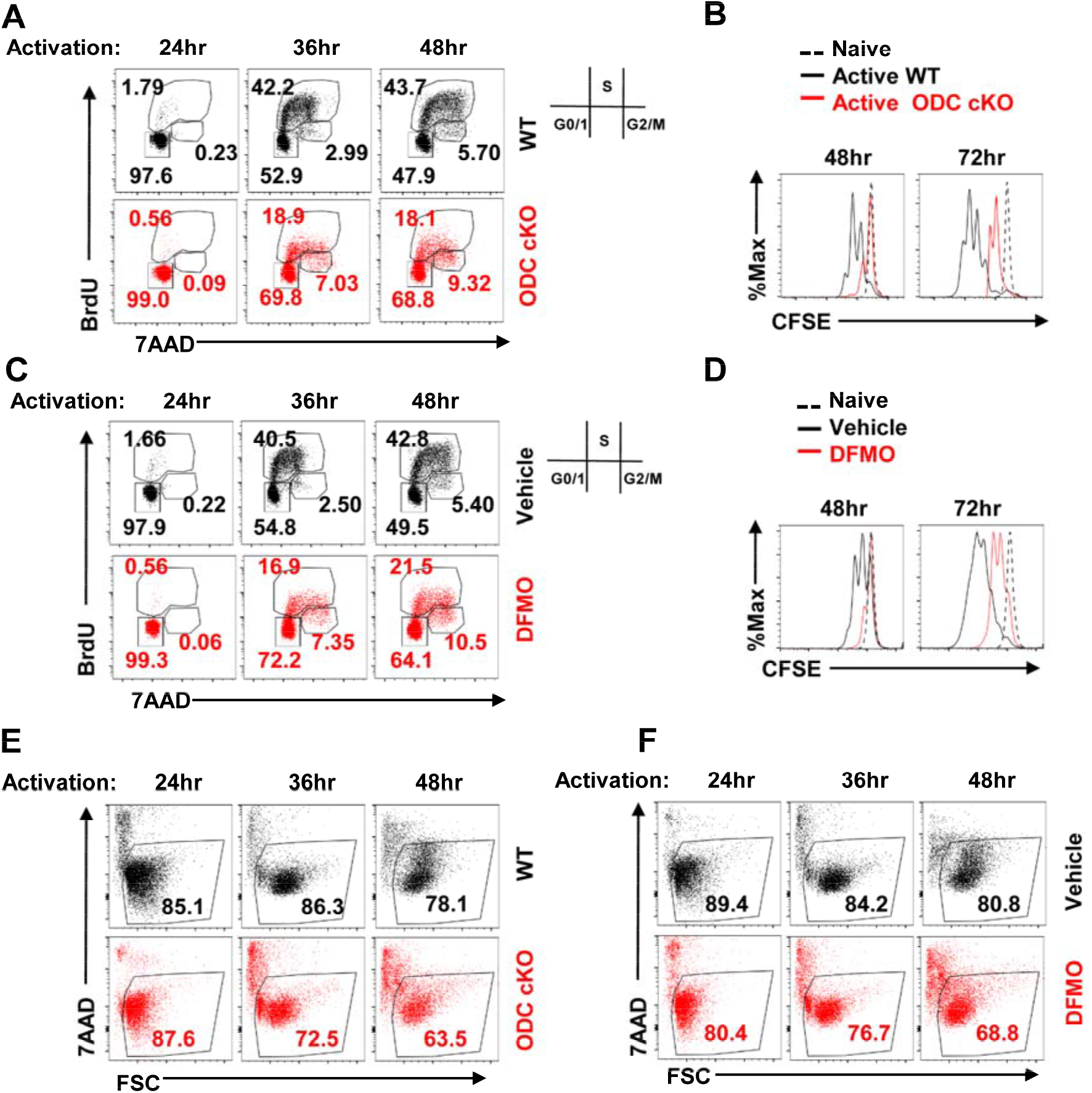
Blockage of de novo polyamine biosynthesis suppresses T cell proliferation and reduces viability *in vitro*. (**A-F**) Naïve T cells isolated from the spleen and lymph nodes of WT or ODC cKO (CD4-Cre, ODC^fl^) mice were activated by plate-bound anti-CD3 and anti-CD28 antibodies with 5 ng/mL IL-2. (**A**), activated WT or ODC cKO T cells were pulsed with BrdU for 1 hour before harvest at indicated time points, then stained with intracellular BrdU antibody and DNA marker 7AAD. Cell cycle profile was analyzed by flow cytometry. The numbers indicate the percentage of cells in each cell cycle stage. (**B**), total T cells isolated from WT or ODC cKO mice were stained with cytoplasmic dye CFSE as indicated. Cell proliferation was evaluated by CFSE dilution with flow cytometry. (**C**), the cell cycle profile was evaluated in active T cells in the presence or absence of 2mM DFMO using BrdU (**D**), cell proliferation profile was analyzed for naïve and activated T cells that received either vehicle or 2mM DFMO treatment. (**E-F**), activated T cells under indicated conditions were analyzed for cell viability using 7AAD uptake which indicates the loss of cell membrane integrity, FSC (forward scatter) reflects cell size change. (**A, C**) representative of 3 independent experiments, (**B, D, E, F**) representative of 6 independent experiments.

### ODC activity is dispensable for T cell proliferation and function *in vivo*

Controlling polyamine homeostasis is crucial for supporting cellular functions in all cell types, and both de novo biosynthesis and salvage by importing extracellular polyamine into the cell tightly regulates cellular polyamine-pool size (*16*–*18*). Without exogenous polyamine supplements in cell culture media, we envisioned that T cells would solely depend on de novo biosynthesis to maintain the intracellular polyamine pool *in vitro*. However, circulating polyamines that are provided by dietary intake and by intestinal microbiota may be a source of exogenous polyamine for T cells *in vivo* (*15*, *41*). To assess the impact of ablating de novo biosynthesis on CD4 T cell proliferation *in vivo*, we employed a well-established competitive homeostatic proliferation assay to determine the ratio and carboxyfluorescein succinimidyl ester (CFSE) dilution pattern of purified *WT(Thy1.1^+^)* or *ODC*^−/−^*(Thy1.2^+^)* CD4^+^ T cells in Rag1-deficient mice. Surprisingly, the ratio between *WT* and *ODC*^−/−^ CD4^+^ T cells was similar before and after adoptive transfer. Additionally, *WT* and *ODC*^−/−^ CD4^+^ T cells display an overlapped CFSE dilution pattern, indicating that the loss of ODC did not affect T cell proliferation *in vivo* (Fig. 2A). Next, we sought to measure antigen-specific, TCR-dependent proliferation of *WT* or *ODC*^−/−^ CD4^+^ T cells. We crossed Thy1.1 and CD4-Cre, ODC^fl^ mice with OT-II transgenic mice to generate *WT(Thy1.1^+^)* and *ODC*^−/−^*(Thy1.2^+^)* donor OT-II strains in CD45.2^+^ background. We then adoptively transferred mixed and CFSE labelled *WT* and *ODC*^−/−^ CD4^+^ T cells into CD45.1^+^ mice that were immunized with chicken ovalbumin 323-339 peptide (OVA_323-339_) in complete Freund’s adjuvant (CFA). After 7 days, we measured the percentage ratio and CFSE dilution pattern of *WT(Thy1.1^+^)* and *ODC*^−/−^*(Thy1.2^+^)* CD4^+^ T cells in popliteal lymph node. Consistent with the homeostatic proliferation results, *WT* and *ODC*^−/−^ OT-II specific CD4^+^ T cells display a comparable antigen-specific proliferation (Fig. 2B). The expansion and balance between pro-inflammatory CD4^+^ T effector (T_eff_) cells determine the pathogenic development of experimental autoimmune encephalomyelitis (EAE), a murine model of multiple sclerosis (MS), which is an inflammatory demyelinating disease of the central nervous system (CNS). We employed this well-characterized system to further interrogate an *in vivo* CD4 T cell response in the absence of polyamine de novo biosynthesis. In line with our homeostatic and antigen-specific proliferation data, neither the genetic deletion of ODC in T cells nor the systemic delivery of DFMO changes the kinetics of pathogenic progression (Fig. 2C and 2D). Together, our data indicates that polyamine salvage from circulation may be able to support T cell proliferation and effector function by compensating for the loss of de novo biosynthesis in the in vivo environment.

**Figure 2.**
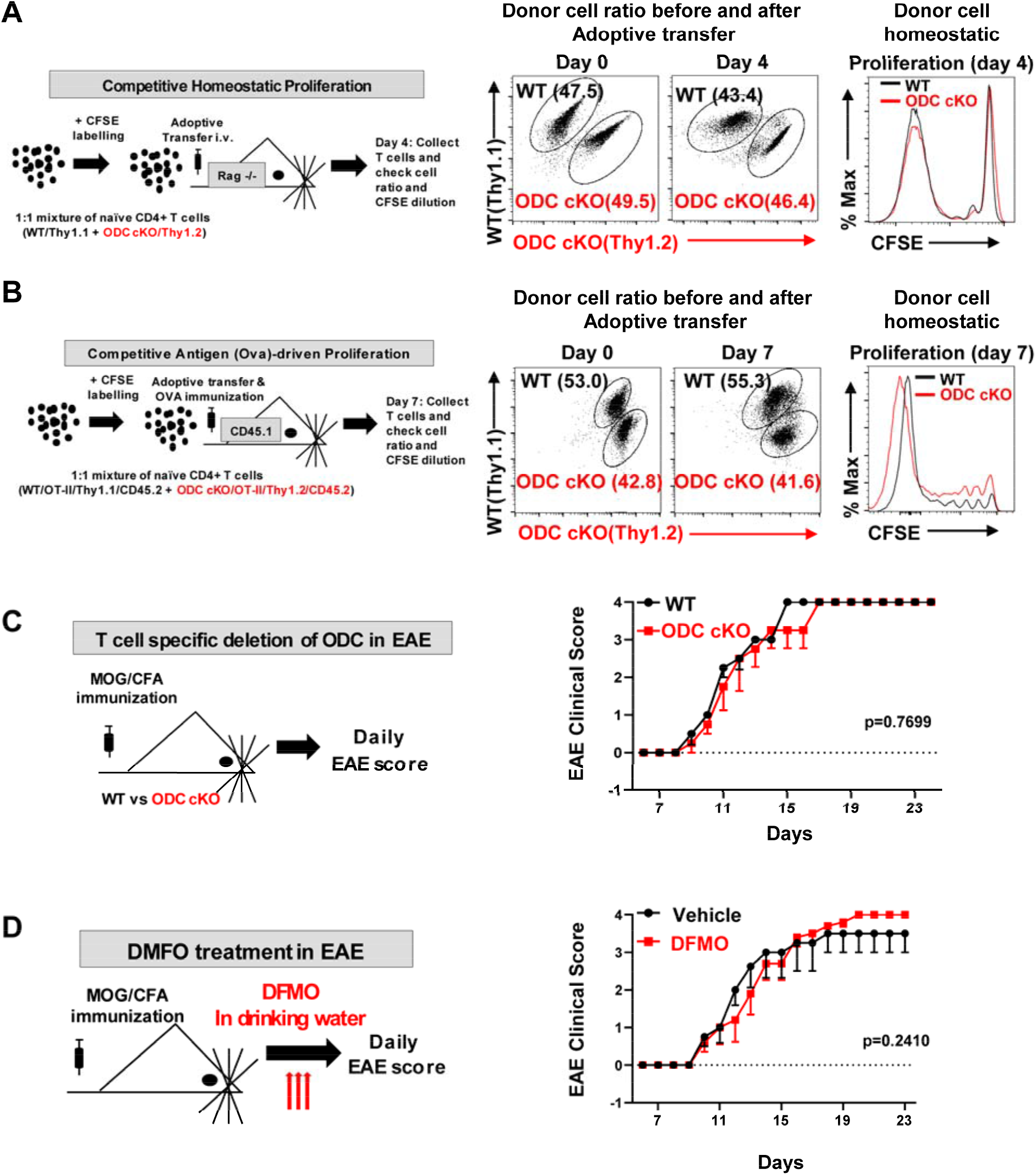
De novo polyamine biosynthesis is dispensable for driving T cell proliferation and function *in vivo*. (**A**), An overview of *in vivo* competitive homeostatic proliferation experimental procedure (left panel). Naïve CD4+ T cells isolated from WT mice (Thy1.1+) and ODC cKO mice (Thy1.2+) were mixed in a 1:1 ratio and stained with CFSE, then adoptively transferred into lymphopenic *Rag*−/− host mice for *in vivo* homeostatic proliferation. The T cell ratio and proliferation were evaluated in WT and ODC cKO cells by Thy1.1/Thy1.2 cell surface staining and CFSE dilution, respectively, by flow cytometry (right panel). (**B**), An overview of *in vivo* competitive OT-II T cell antigen (ovalbumin/OVA) driven proliferation experimental procedure (left panel). Naïve CD4+ OT-II T cells isolated from WT OT-II mice (Thy1.1+, CD45.2+) and ODC cKO OT-II mice (Thy1.2^+^, CD45.2+) were mixed in a 1:1 ratio and labeled with CFSE, then adoptively transferred into CD45.1+ host mice, which were immunized in the hock with OVA peptides. After 7 days, popliteal draining lymph nodes were collected, and donor cell ratio and proliferation were analyzed by flow cytometry. Data are representative of 2 independent experiments. (**C**), An overview of induced experimental autoimmune encephalomyelitis (EAE) experimental procedure. Mice with indicated genotypes were immunized with MOG/CFA to induce EAE, and clinical scores were evaluated daily. (**D**), WT mice were immunized with MOG/CFA, and treated with 1% DFMO in drinking water or regular drinking water (vehicle) from day 0 throughout the experiment. Clinical scores were evaluated daily. (**C, D**) EAE data indicate mean ± SEM. N=5. (**A, C, D**) representative of 3 independent experiments.

### Polyamine salvage compensates for the loss of biosynthesis activity *in vitro*

Next, we investigated the role of polyamine salvage (uptake) in regulating polyamine homeostasis in T cell. We measured polyamine uptake activity using radioactive-labelled putrescine (^14^C-Putrescine) in naïve, active *WT* and *ODC*^−/−^ T cells. Active T cells displayed higher polyamine uptake activity than naïve T cells (Fig. 3A). Importantly, ablation of ODC induces a compensatory increase in polyamine uptake (Fig. 3B). Next, we asked if polyamine uptake is sufficient to maintain polyamine homeostasis and support T cell proliferation in the absence of ODC activity. While genetic deletion or pharmacologic inhibition of ODC significantly delayed cell cycle progression from G0/G1 to the S phase and suppressed overall T cell proliferation, exogenous polyamine supplement could restore the cell cycle progression, proliferation and viability in DMFO-treated and *ODC*^−/−^ CD4^+^ T cells *in vitro* (Fig. 3C-3F and Fig. S3A-S3B). We then sought to employ a pharmacologic approach to block polyamine uptake and assessed the role of polyamine uptake in regulating polyamine homeostasis. AMXT 1501 (AMXT) is a novel lipophilic polyamine mimetic that potently blocks polyamine uptake in the low nanomolar concentration (*42*). The combination of AMXT and DFMO could effectively deplete the polyamine pool in tumor cells and suppress the growth of tumors in various animal models (*43*–*45*). These promising preclinical studies led to a recently opened Phase I clinical trial in solid tumors (<u>NCT03077477</u>). Similar to the genetic data (Fig. 3B), DFMO treatment induces a compensatory increase in polyamine uptake, which can be blocked by AMXT 1501 (Fig. 3G). In addition, AMXT 1501 could significantly suppresses exogenous polyamine-mediated cell proliferation and viability in *ODC*^−/−^, but not *WT* CD4^+^ T cells (Fig. 3H and S3C), indicating that either polyamine salvage or biosynthesis is sufficient to maintain polyamine homeostasis in T cells. Supporting this idea, AMXT 1501 treatment alone failed to suppress T cell homeostatic proliferation or antigen-specific proliferation (Fig. S4A and S4B). These results, together with the results described above, suggest that the salvage pathway and the de novo biosynthesis pathway can compensate for the loss of each other, representing a layer of metabolic plasticity engaged by T cells to maintain polyamine homeostasis.

**Figure 3.**
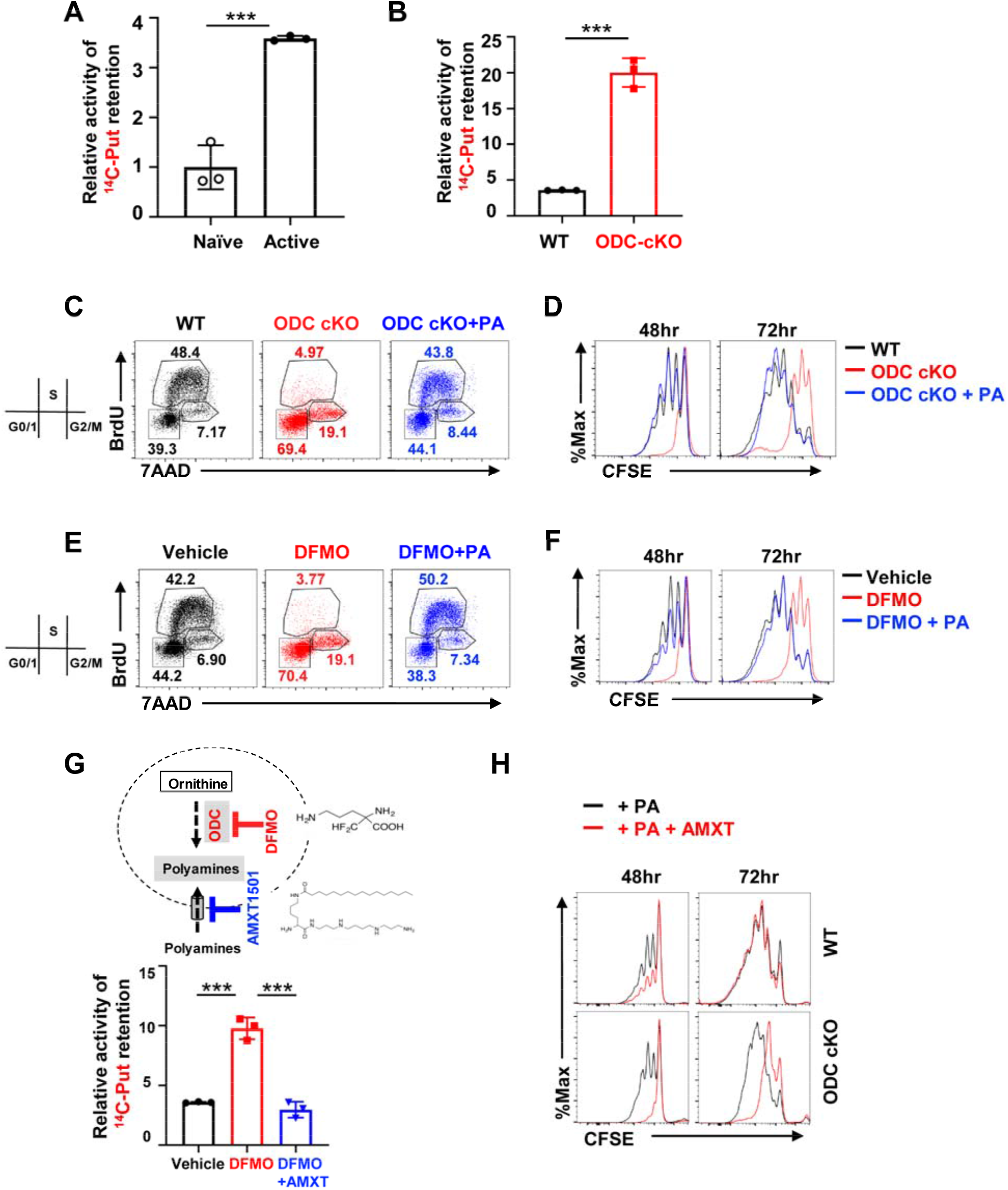
Ablation of de novo polyamine biosynthesis renders T cells dependent on polyamine uptake that can be blocked by AMXT1501. (**A**), Relative putrescine uptake of naïve and activated T cells was determined by the cellular retention of ^14^C labeled putrescine. (**B**), relative putrescine uptake of WT and ODC cKO cells after 24 hours activation. (**C-F**), naïve T cells isolated from WT or ODC cKO mice were activated by plate-bound antibodies in presence of 5 ng/mL IL-2. (**C**), Cell cycle profile was evaluated in WT, ODC cKO, and ODC cKO groups supplemented with 3μM polyamine mixture (putrescine 1 μM, spermidine 1 μM, spermine 1 μM) using BrdU after 48hour of T cell activation. The numbers indicate the percentage of cells in each cell cycle stage. (**D**), Cell proliferation was evaluated as indicated condition by CFSE dilution. (**E**), isolated T cells with control vehicle (PBS as solvent), 2 mM DFMO (ODC inhibitor), or 2 mM DFMO with 3 μM PA mixture treatment were activated, BrdU pulsed, and analyzed for cell cycle profile as described in **C**. (**F**), cell proliferation profile at indicated time point was analyzed for activated T cells in presence and absence of, 2 mM DFMO, or 2 mM DFMO plus 3 μM PA mixture as described in **D**. (**G**), overview of cellular polyamine homeostasis through biosynthesis and uptake as well as the pharmacological inhibitors’ function of depleting intracellular polyamine pool (top panel). The effect of DFMO alone or combined with polyamine uptake inhibitor AMXT1501 (1 μM) on polyamine uptake in 24-hour activated T cells was evaluated as described in **A.** (**H**), Cell proliferation profile was evaluated for indicated condition by CFSE dilution. (**A, B, G**) representative of 4 independent experiments. (**C, D, E, F, H**) representative of 3 independent experiments. Bar graphs indicate mean ± SD.

### Simultaneously blocking polyamine salvage and biosynthesis suppresses T cell proliferation *in vivo*

Given that either genetically ablating ODC or pharmacologically blocking polyamine uptake failed to impact T cell proliferation and function *in vivo*, we sought to assess the impacts of simultaneously blocking polyamine salvage and biosynthesis on T cells *in vivo*. We employed competitive homeostatic proliferation and antigen-specific T cell proliferation assays to determine the proliferation of *WT* and *ODC*^−/−^ donor CD4^+^ T cells in recipient mice, which were treated with vehicle or AMXT 1501 during the course of experiment. Donor *ODC*^−/−^ CD4^+^ T cells recovered from AMXT 1501 treated animals but not from vehicle treated animals displayed reduced percentage and delayed proliferation compared with competitive *WT* CD4^+^ T cells (Fig. 4A and 4B). We then sought to assess the impacts of simultaneously blocking polyamine salvage and biosynthesis on T cells in the EAE model. The genetic deletion of ODC in T cells failed to cause any significant changes in EAE pathogenic progression. Animals that were treated with AMXT 1501 displayed a delayed disease onset initially, but eventually proceeded with pathologic development and reached the endpoint. Importantly, the combination of AMXT 1501 with genetic deletion of ODC in T cells conferred full protection against EAE pathogenic progression (Fig. 4C). ODC inhibitor DFMO is an FDA-approved medicine for hirsutism and African sleeping sickness and has been widely tested as a chemopreventive and chemotherapeutic agent against solid tumors (*46*–*51*). Similar to the genetic data, DFMO alone failed to suppress EAE pathogenic progression (Fig. 4D). Although AMXT alone was sufficient to delay EAE onset moderately, it failed to protect animals from reaching the endpoint (Fig. 4D). We envision that the combination of DFMO and AMXT may be sufficient to deplete T cell polyamine pool, and consequently suppress T cell proliferation and effector function *in vivo*. Supporting this idea, the combination of AMXT and DMFO, but not single treatments, conferred full protection against EAE pathogenic progression (Fig. 4D). Inflammatory T_H_1, T_H_17, and FoxP3-expressing regulatory T cells (T_reg_) are closely related to CD4 T cell subsets but with distinct functions. The balance between pro-inflammatory CD4^+^ T_eff_ cells and T_reg_ cells determines the pathogenic development of EAE. Next, we examined the polyamine’s role in CD4 T_eff_ cell differentiation *in vitro*. Without exogenous polyamine supplemented in cell culture media, maintenance of the intracellular polyamine pool depends solely on ODC-mediated polyamine biosynthesis. Remarkably, intracellular polyamine depletion resulting from ODC deficiency inhibited pro-inflammatory T_H_1 and T_H_17 cell differentiation while enhancing anti-inflammatory iT_reg_ cell differentiation in vitro (Fig. S5). Together, our results indicate that the polyamine blocking approach, via ablation of salvage and biosynthesis pathways, suppresses T cell proliferation and may be a potential new therapy for treating inflammatory and autoimmune disease.

**Figure 4.**
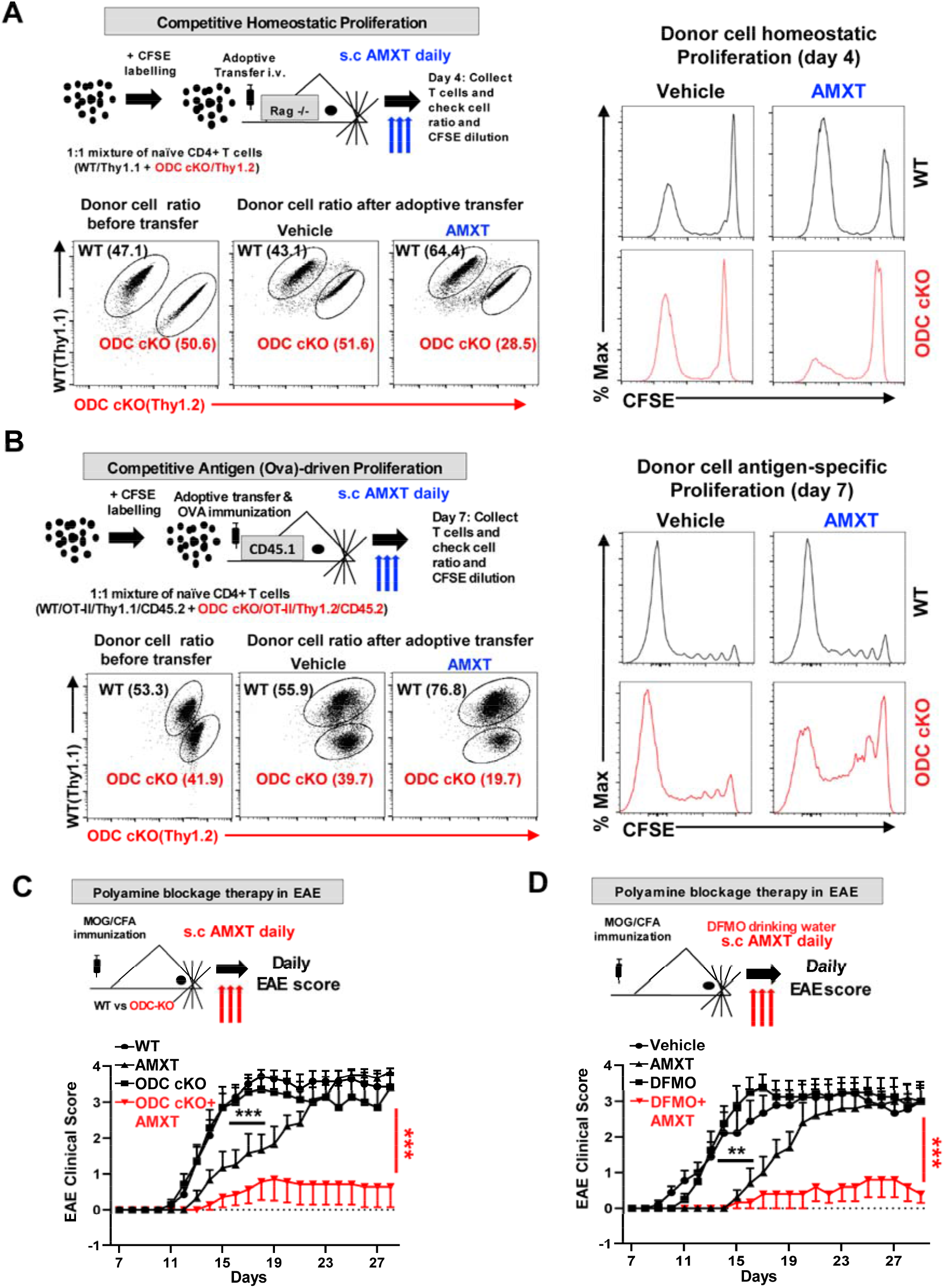
Simultaneous blockage of polyamine uptake and de novo biosynthesis suppresses T cell proliferation and function *in vivo*. (**A**), overview of *in vivo* competitive homeostatic proliferation experimental procedure (top panel). Naïve CD4+ T cells isolated from WT mice (Thy1.1+) and ODC cKO mice (Thy1.2+) were mixed in a 1:1 ratio and stained with CFSE, then adoptively transferred into lymphopenic *Rag*^−/−^ host mice for *in vivo* homeostatic proliferation. Host mice were randomly divided into two groups and treated with vehicle (PBS as control) or AMXT1501 (3mg/kg/day s.c.) for 4 days. Peripheral lymph nodes were collected, and the percentages of WT and ODC cKO cells as well as cell proliferation were analyzed by Thy1.1/Thy1.2 cell surface staining and CFSE dilution by flow cytometry, respectively. (**B**), An overview of *in vivo* competitive OT-II T cell antigen (ovalbumin/OVA) driven proliferation experimental procedure (top panel). Naïve CD4+ OT-II T cells isolated from WT OT-II mice (Thy1.1+, CD45.2+) and ODC cKO OT-II mice (Thy1.2+, CD45.2+) were mixed in a 1:1 ratio and labeled with CFSE, then adoptively transferred into CD45.1+ host mice, which were then immunized in the hock with OVA peptides and treated with vehicle (PBS as control) or AMXT1501 (3mg/kg/day s.c.). After 7 days of *in vivo* proliferation, draining lymph nodes were collected, percentage and proliferation of donor cells were analyzed by flow cytometry. (A-B) representative of two independent experiments. (**C**), An overview of induced experimental autoimmune encephalomyelitis (EAE) (top panel). Mice with indicated genotypes were immunized with MOG/CFA to induce EAE. Each sub-group from WT and ODC cKO mice were treated with polyamine uptake inhibitor AMXT1501 (3mg/kg/day s.c.), and clinical scores were evaluated for 4 groups daily. Data represents 2 independent experiments. (**D**), The schematic diagram of DFMO and AMXT treatment in EAE model. WT mice were immunized with MOG/CFA, randomized into four groups for the treatment with vehicle (PBS as control), AMXT1501(3mg/kg/day s.c.), DFMO (1% in drinking water), and DFMO (1% in drinking water) + AMXT1501(3mg/kg/day s.c.), respectively, throughout the experiment. Clinical scores were evaluated daily. Data represents 2 independent experiments. EAE data indicate mean ± SEM. N among 5 to 10.

### Amino acid catabolism provides carbon sources for polyamine biosynthesis

Dietary intake and intestinal microbiota metabolism are major sources of circulating polyamine (*15*, *41*). We reasoned that an understanding of carbon sources that drive polyamine biosynthesis in T cells would enable the development of nutritional approaches for effectively controlling polyamine homeostasis in T cells. While arginine catabolism is integrated into the urea cycle to detoxify ammonia and provides the precursor for polyamine biosynthesis, we previously reported that glutamine-derived carbon could be funneled into polyamine biosynthesis through ornithine in T cells (*24*). In addition, proline can also provide carbon for polyamine biosynthesis in the placenta and in plants (*13*, *52*). To determine to what extent these three amino acids contribute to polyamine biosynthesis, we applied the stable isotope of carbon-13 (^13^C) labeling and mass spectrometry approach. We supplied ^13^C_6_-Arginine, ^13^C_5_-Glutamine, or ^13^C_5_-Proline as metabolic tracers in T cell culture media and then followed ^13^C incorporation into individual metabolites (Fig. 5A). While not all desired metabolites were detected in our experiment due to technical limitation and a portion of proline was produced through de novo biosynthesis, our results clearly demonstrated that ^13^C_5_-Proline only contributes a minimal amount of ^13^C_5_ isotopologue of ornithine and polyamine (Fig. 5B). In contrast, ^13^C_6_-Arginine and ^13^C_5_-Glutamine contribute 50% and 40% of ^13^C_5_-Ornithine, respectively (Fig. 5B). Importantly, ^13^C_6_-Arginine and ^13^C_5_-Glutamine contribute around 80% and 20% of ^13^C_4_ isotopologues of polyamine (putrescine and spermidine generated via decarboxylation of ornithine), respectively (Fig. 5B). Thus, we conclude that arginine is a major carbon donor and glutamine is a minor carbon donor for supporting polyamine biosynthesis in T cells *in vitro*.

**Figure 5.**
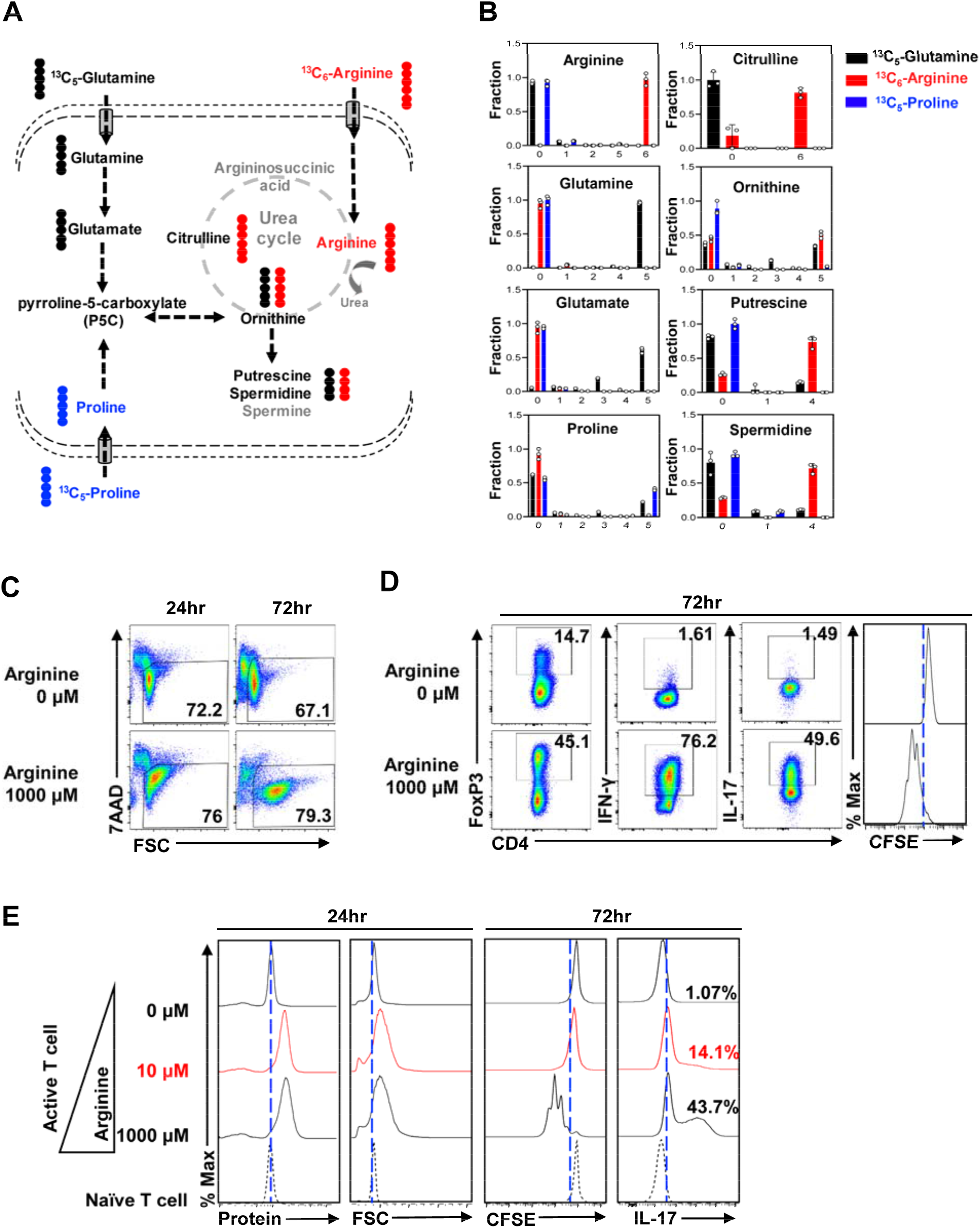
Arginine catabolism supports polyamine biosynthesis and T cell proliferation. (**A**), diagram of ^13^C_5_-Glutamine, ^13^C_6_-Arginine, and ^13^C_5_-Proline entering the intracellular polyamine biosynthesis pathways. (**B**), Naive T cells isolated from spleen and lymph nodes of WT mice were activated by plate-bound method in Gln/Arg/Pro triple-free media supplemented with ^13^C_5_-Glutamine or ^13^C_6_-Arginine, or ^13^C_5_-Proline respectively for 36 hours, extracted, and analyzed for indicated metabolites using CE-TOFMS as described in methods. (**C**), T cell viability in the indicated conditional media at designated time point was determined by 7AAD uptake. (**D**), naïve CD4+ T cells isolated from WT mice were labeled with CFSE, activated, and differentiated in arginine-free RPMI1640 or high-level arginine supplemented (1000μM, which comparable with complete RPMI 1640) media for 72 hours. The indicated proteins IFN-ɣ (T_H_1), IL-17 (T_H_17), and FoxP3 (iT_reg_) were quantified by intracellular staining following PMA and ionomycin stimulation. Cell proliferation in arginine-free or high-level arginine supplemented media was determined by CFSE dilution. (**E**), isolated T cells were either maintained in culture media containing 5 ng/mL IL-7 as naïve T cells or activated in the arginine-free conditional media with 5 ng/mL IL-2 and increased concentrations of arginine (0 μM, 10 μM, and 1000 μM), respectively. T cell protein synthesis levels (FITC staining) and cell size (FSC) were evaluated at 24 hours. Cell proliferation (CFSE) and its ability to polarize into T_H_17 (IL-17) were evaluated at 72 hours. (**B, E**) representative of 2 independent experiments. (**C**) representative of 5 independent experiments. (**D**) representative of 3 independent experiments. Bar graphs indicate mean ± SD.

### Arginine catabolism drives T cell proliferation partially through supporting polyamine biosynthesis

While arginine is generally considered a non-essential amino acid, T cells are arginine auxotrophic *in vitro* (*29*, *53*, *54*). Arginine is required to maintain CD4^+^ T cells viability, driving proliferative and proinflammatory lineage (T_H_17) differentiation (Fig. 5C and 5D). While the amount of most amino acids is in the low micromolar (μM) range, the concentration of arginine is in the millimolar (mM) range in cell culture media. We envisioned that, in addition to a general requirement of arginine for protein synthesis, arginine might also support T cell proliferation and differentiation through polyamine biosynthesis. To test this idea, we titrated down the amount of arginine in cell culture media and found that 10 μM of arginine was sufficient to support protein synthesis and the growth of cell mass at the early time point after T cell activation but failed to maintain cell viability and drive proliferation later (Fig. S6A and 5E). We envision that arginine supports protein synthesis and polyamine biosynthesis, both of which are required to drive T cell proliferation. Supporting this idea, we have shown that a low level of arginine (10 μM) in culture media is sufficient to support protein synthesis, but not proliferation (Fig. 5E). Similar to what we found in polyamine depletion condition (Fig. S5), low level of arginine in culture media reduces T_H_17 differentiation (Fig. 5E). However, polyamine supplements can partially restore T cell proliferation and T_H_17 differentiation with a low level of arginine (10 μM) (Fig. S6C). In contrast, polyamine supplements failed to restore T cell proliferation and differentiation under arginine starvation condition (0 μM) (Fig. S6B). Collectively, our data indicate that T cell activation engages a metabolic axis that connects arginine catabolism to polyamine de novo biosynthesis.

## Discussion

Fast-growing pathogens impose selective pressures on the metabolic fitness and metabolic plasticity of host immune cells, which are necessary for immune cells to maintain homeostasis while remaining ready to mount rapid responses under diverse metabolic and immune conditions (*3*, *55*–*58*). A robust T cell-mediated adaptive immune response exhibits high and dynamic metabolic demands in T cells, which is accommodated through fine-tuned regulation on both the central carbon metabolic pathways, and peripheral metabolic pathways including polyamine metabolism (*38*, *59*–*65*). The standard formulation of cell culture media, which does not fully recapitulate the physiological metabolite composition in the plasma and tissue microenvironment, often leads to a significant metabolic adaptation of growing cells *in vitro*. Metabolic phenotypes of cells growing *in vivo* may further differ from cells growing *in vitro* as a result of other environmental factors including oxygen level, cellular competition and cooperation, and the biophysical properties of the extracellular matrix (*66*–*71*). Similarly, T cell metabolism can respond and adapt to environmental nutrient levels (*3*, *37*). Our studies revealed T cells’ capacity to engage in both de novo biosynthetic and salvage pathways to fine-tune polyamine homeostasis, which is required to maximize metabolic fitness and optimize CD4 T_eff_ cell proliferation and differentiation. Such metabolic plasticity is likely to be crucial for T cells’ ability to elicit robust immune responses in different tissue contexts.

Glutamine and arginine are two non-essential amino acids that not only fulfill the general requirements for protein synthesis, but also connect central carbon metabolism to a variety of biosynthetic pathways to produce specialized metabolites (*22*, *72*–*75*). Glutamine catabolism funnels the anaplerotic flux of carbon into the TCA cycle and also provides sources of nitrogen and carbon to support biosynthesis of nonessential amino acids, lipids, nucleotides, glutathione and polyamines in T cells (*24*, *38*, *76*–*79*). Similarly, arginine is critical for maintaining T cells viability, driving proliferative and effector functions (*25*, *29*, *54*, *80*). We found that polyamine supplement could partially relieve T cell’s dependence on arginine, indicating that a key role of arginine catabolism in T cells is to support polyamine biosynthesis. Consistently, arginine serves as a major donor of carbon in polyamine biosynthesis, while glutamine only plays a minor role in funneling carbon into polyamine in the presence of arginine in vitro. Interestingly, these two amino acids contribute a comparable portion of carbon to ornithine, the precursor of polyamine. This finding is in line with previous studies showing that arginine-derived ornithine is not the only source for endogenous ornithine in most mammalian tissues (*81*, *82*). In addition, our findings may implicate ornithine as an important metabolic node representing a key branch point in both glutamine and arginine catabolic pathways. Ornithine can be committed towards the urea cycle, proline biosynthesis or de novo synthesis of polyamine. The production of ornithine from two different metabolic precursors, glutamine and arginine, also enables fine-tuned coordination between the metabolic flux shunted towards polyamine synthesis and the metabolic flux shunted towards other specialized metabolites. Consistent with this idea, the overall high consumption rate of glutamine and arginine may provide a sensitive and precise regulation on intermediate metabolites that can be committed toward several metabolic branches, hence permitting rapid responses to meet the metabolic demands of cell growth and cytokine production.

Active T cells and other highly proliferative cells such as tumor cells share metabolic characteristics and are strictly dependent on the catabolism of glucose and glutamine through the central carbon metabolic pathways (*83*–*90*). Similarly, elevated levels of polyamine are associated with T cell activation and cell transformation (*24*, *91*, *92*). Importantly, pharmacologic or genetic targeting of ODC, a transcriptional target of proto-oncogene MYC, could delay the development and the progression of MYC-driven tumors in mice (*93*, *94*). While pharmacologic agents targeting polyamines such as the ODC inhibitor DFMO has generally low toxicity, DFMO yields only very marginal therapeutic benefits in cancer patients as a single-agent therapy in clinical trials (*50*). Clearly, cancer cells are capable of importing polyamine from circulation to overcome the effect of DFMO. Supporting this idea, the simultaneous blockage of polyamine biosynthesis and uptake has generated promising results in tumor preclinical animal models (*43*, *45*). Mammalian cells can uptake polyamine through endocytosis and membrane transport system mediated by several solute carrier transporters (*95*–*99*). AMXT 1501 is a novel lipophilic polyamine mimetic that potently blocks polyamine uptake through competing with polyamine (*42*). The combination of AMXT 1501 and DFMO could effectively deplete the polyamine pool in tumor cells and suppress the growth of tumors in various animal models (*43*–*45*). These promising preclinical studies led to a recently opened Phase I clinical trial in solid tumors (<u>NCT03077477</u>). Similarly, we have shown that ODC^−/−^ CD4^+^ T cells proliferate and function normal *in vivo*. However, concurrent treatment of ODC^−/−^ CD4^+^ T cells with AMXT 1501 abrogate their proliferation and inflammatory function *in vivo*. Moreover, the combination of AMXT 1501 and DFMO could confer full protection to mice against pathogenic development of EAE. Thus, polyamine blocking strategy via simultaneous blockade of polyamine biosynthesis and salvage may present a promising and novel therapy for treating inflammatory and autoimmune diseases.

## Materials and Methods

### Mice

C57BL/6NHsd (WT) mice, Rag−/− mice, OTII mice, Thy 1.1+ mice, and CD45.1+ mice were obtained from Jackson laboratory (Bar Harbor, ME). CD4-Cre, ODC^fl^ (ODC cKO) mice with C57BL/6 background were produced by FRT-LacZ; ODC^fl^ mice (European Mouse Mutant Archive) crossed with the FLP knock-in mouse strain to remove the LacZ-reporter allele, and the generated mouse strain containing the conditional allele (ODC^fl^) was further crossed with the CD4-Cre strain. For experiments, gender and age matched mice around 6-12 weeks of age kept in specific pathogen-free conditions were used. All animal experiment protocols were approved by the Institutional Animal Care and Use Committee of Abigail Wexner Research Institute at Nationwide Children’s Hospital (IACUC; protocol number AR13-00055).

### Flow cytometry

For analysis of surface markers, cells were stained in PBS containing 2% (w/v) BSA and the appropriate antibodies from Biolegend. Foxp3 expression was performed using the Foxp3 staining kit from eBioscience. For intracellular cytokine IFN-ɣ and IL-17A staining, T cells were stimulated for 4 hours with phorbol 12-myristate 13-acetate (PMA) and ionomycin in the presence of monensin before being stained with CD4 antibody. Cells were then fixed and permeabilized using Foxp3 Fixation/Permeabilization solution according to the manufacturer’s instructions (eBioscience™). Cell total protein level was assessed by intracellular FITC (Fisher scientific) staining. Cell proliferation was assessed by CFSE staining per the manufacturer’s instructions (Invitrogen). Cell viability was assessed by 7-AAD staining per the manufacturer’s instructions (Biolegend). Flow cytometry data were acquired on Novocyte (ACEA Biosciences) and were analyzed with FlowJo software (TreeStar).

### Cell Culture

For *in vitro* culture, total T cells or naïve CD4+ T cells were isolated from mouse spleen and lymph nodes using MojoSort mouse CD3/CD4 naïve T cell Isolation Kit (Biolegend) following the manufacturer’s instructions. For all *in vitro* cell culture, unless indicated separately, complete RPMI-1640 medium (containing 10% (v/v) heat-inactivated dialyzed fetal bovine serum (DFBS), 2 mM L-glutamine, 0.05 mM 2-mercaptoethanol, 100 units/mL penicillin, and 100 μg/mL streptomycin) was used. DFBS was made by dialyzing against 100 volumes of PBS (five changes in three days) using Slide-ALyzerTM G2 dialysis cassettes with cut-through MW size 2K (ThermoFisher Scientific) at 4 ⁰C to remove any potential polyamines in the FBS. For the activation assay, freshly isolated total T cells were either maintained in culture media with 5 ng/mL IL-7 for resting state or were activated with 5 ng/mL IL-2 and plate-bound anti-CD3 (clone 145-2C11) and anti-CD28 (clone 37.51). Plates were pre-coated with 2 μg/mL antibodies overnight at 4°C. Cells were seeded as 1 × 10^6^ cells/mL and cultured in RPMI 1640 media at 37 °C in 5% CO_2_. For CFSE dilution analysis, cells were pre-incubated for 10 min in 4 μM CFSE (Invitrogen) in PBS plus 5% FBS before culture. For induced T_reg_ cell differentiation, 0.5 ×10^6^ naïve CD4+ T cells were cultured with 200 U/mL IL-2, and 5 ng/mL TGF-β in 0.5 mL RPMI-1640 media in a 48-well tissue culture plate that was pre-coated with 10 μg/mL anti-CD3 and 10 μg/mL anti-CD28 overnight at 4°C. For T_H_17 differentiation condition, 0.5 ×10^6^ naïve CD4+ T cells were seeded in each well pre-coated with 10 μg/mL anti-CD3 and 10 μg/mL anti-CD28 overnight at 4°C and cultured with 8 μg/mL anti–IL-2, 8 μg/mL anti–IL-4, 8 μg/mL anti–IFN-γ, 2 ng/mL TGF-β, and 20-50 ng/mL IL-6 in 0.5mL RPMI-1640 media in a 48-well tissue culture plate. For T_H_1 differentiation condition, 0.5 ×10^6^ naïve CD4+ T cells were seeded in a well pre-coated with 10 μg/mL anti-CD3 and 10 μg/mL anti-CD28 overnight at 4°C and cultured with 5 ng/mL IL-12, 10 μg/mL anti–IL-4, and 200 U/mL IL-2 in 0.5mL RPMI-1640 media in a 48-well tissue culture plate for 72 hours. For invitro, cell culture experiments, ODC inhibitor DFMO (Carbosynth), 2 mM, polyamine uptake inhibitor AMXT 1501 (Aminex Therapeutics), 1 μM was used for polyamine mixture, unless specifically indicated, a 3 μM polyamine mixture (putrescine 1μM, spermidine 1μM, spermine 1μM) was used for cell culture. In order to prevent diamine oxidase in the FBS from breaking down the polyamine supplement in cell culture, 0.2 mM aminoguanidine (Sigma) was added to all polyamine supplemented groups.

### Cell cycle analysis

Upon *in vitro* activation, T cell cycle analysis was performed using Phase-Flow Alexa Fluro 647 BrdU Kit (Biolegend) per the manufacturer’s instructions. Briefly, T cells were pulsed with 10 μg/mL BrdU for 1 hour before being processed with surface staining, fixation, and permeabilization. BrdU incorporated into the DNA during S phase were recognized by intracellular BrdU antibody staining, and total DNA content was used to differentiate G1 and G2 stages by 7AAD labeling.

### qPCR

Total RNA was isolated using the RNeasy Mini Kit (Qiagen) and was reverse transcribed using random hexamers and M-MLV Reverse Transcriptase (Invitrogen). SYBR green-based quantitative RT-PCR was performed using the Applied Biosystems 7900 Real Time PCR System. The relative gene expression was determined by the comparative *C*_T_ method, also referred to as the 2^−ΔΔ*C*_T_^ method. The data were presented as the fold change in gene expression normalized to an internal reference gene (beta2-microglobulin) and relative to the control (the first sample in the group). Fold change=2^−ΔΔ*C*_T_^=[(*C*_Tgene of interst_− *C*_Tinternal reference_)]sample A−[(*C*_Tgene of interst_− *C*_Tinternal reference_)]sample Samples for each experimental condition were run in triplicated PCR reactions. Primer sequences were obtained from Primer Bank.

### Putrescine uptake assay

2×10^6^ T cells were suspended in 200ul PBS containing permeant putrescine (100 μM) and [1,4- ^14^C]-putrescine dihydrochloride (0.2 uCi, ARC 0245) and incubated at 37°C for 10 minutes (within the established linear phase of uptake). The reaction was stopped by loading all the transport mixture onto a discontinuous gradient of bromododecane and perchloric acid/sucrose and then centrifuged at 14, 000 ×*g* for 90 seconds. The discontinuous gradient was prepared by overlaying 1-Bromododecane (800 μL; Sigma-Aldrich) above 100 μL of 20% perchloric acid (Sigma-Aldrich)/8% sucrose solution in 1.5-ml microfuge tube. The samples were snap-frozen in an ethanol-dry ice bath. The bottom part of microfuge tubes containing T cell lysate in perchloric acid-sucrose was cut by a microfuge cutter, washed with 300 μL of 0.5%SDS-1%TritonX100, and transferred into scintillation vials. 10 mL scintillation cocktail was then added to each vial, and the radioactivity was then quantitated by liquid scintillation spectrometry.

### Adoptive cell transfer and *in vivo* proliferation

For homeostatic proliferation in lymphopenic *Rag*^−/−^ mice, naïve CD4+ T cells isolated from donor mice using naïve CD4+ mouse T cell isolation kit (Biolegend) were labeled with CFSE. Approximately 1×10^7^ cells in 150 μL PBS were transferred via retro-orbital venous injection into 6-8 week-old gender-matched host mice. Mice were sacrificed after 4 days and lymph nodes were extracted from host mice, then processed for surface staining and flow analysis.

For antigen driven proliferation using OTII mice: naïve CD4+ T cells isolated from OTII/CD45.2 TCR transgenic donor mice using naïve CD4+ mouse T cell isolation kit (Biolegend) were labeled with CFSE. Approximately 1×10^7^ cells in 150 μL PBS were transferred via retro-orbital venous injection into 6-8 week-old gender-matched WT/CD45.1 host mice. Host mice were immunized subcutaneously in the hock area (50 μL each site) in both legs with 1 mg/mL OVA^323-339^ peptide (InvivoGen) emulsified with CFA (InvivoGen). After 7 days of antigen-driven proliferation, lymph organs were extracted from host mice then processed for surface staining and flow analysis.

### Experimental Autoimmune Encephalomyelitis (EAE)

For induced EAE, mice were immunized subcutaneously with 100 μg of myelin oligodendrocyte glycoprotein (MOG)_35–55_ peptide emulsified in complete freund adjuvant (CFA), which was made from IFA(Difco) plus mycobacterium tuberculosis (Difco). Mice were i.p. injected with 200 ng of pertussis toxin (List Biological,#181) on the day of immunization and 2 days later. For the DFMO (Carbosynth) treated group, mice were fed with 1% DFMO in drinking water throughout the experiment. DFMO water was replenished every five days. For the AMXT 1501 (Aminex Therapeutics) treated group, mice received 3mg/kg of AMXT 1501 subcutaneous daily throughout the experiment. The mice were observed daily for clinical signs and scored as described below.

|  |  |
| --- | --- |
| Score 1 | Limp tail. When the mouse is picked up by the tail, the whole tail drapes over your finger, instead of being erect. |
| Score 2 | Limp tail and weakness of hind legs. When mouse is picked up by tail, legs are not spread apart, but held closer together. When the mouse is observed when walking, it has a clearly apparent wobbly walk. |
| Score 3 | Limp tail and complete paralysis of hind legs (most common).<br>OR Limp tail with paralysis of one front and one hind legs. |
| Score 4 | Limp tail, complete hind leg and partial front leg paralysis.<br>Mouse is minimally moving around the cage but appears alert and feeding.<br>Usually, euthanasia is recommended |

### CE-QqQ/TOFMS analysis

Total mouse T cells with 75-80% CD3 positivity were isolated from WT or ODC cKO mice. Isolated cells were activated with plate-bound anti-CD3 (2 μg/mL) and anti-CD28 (2 μg/mL) antibodies with IL-2 (5 ng/mL), and were cultured in 6-well plates for 36 hours. Activated cells (around 1.5×10^7^ cells/sample) was used for the extraction of intracellular metabolites. The cells were collected by centrifugation (300 ×*g* at 4°C for 5 min) and washed twice with 5% mannitol solution (10 mL first and then 2 mL). The cells were then treated with 800 μL of methanol and vortexed for 30 s in order to inactivate enzymes. Next, the cell extract was treated with 550 μL of Milli-Q water containing internal standards (H3304-1002, Human Metabolome Technologies, inc., Tsuruoka, Japan) and vortexed for 30 s. The extract was obtained and centrifuged at 2,300 ×*g* and 4°C for 5 min, and then 700 μL of upper aqueous layer was centrifugally filtered through a Millipore 5-kDa cutoff filter at 9,100 ×*g* and 4°C for 180 min to remove proteins. The filtrate was centrifugally concentrated and re-suspended in 50 μL of Milli-Q water for CE-MS analysis.

Cationic compounds were measured in the positive mode of CE-TOFMS and anionic compounds were measured in the positive and negative modes of CE-MS/MS according to the methods developed by Soga, *et al* [PMID:10740865, PMID:12038746, PMID:14582645].

Peaks detected by CE-TOFMS and CE-MS/MS were extracted using an automatic integration software (MasterHands, Keio University, Tsuruoka, Japan [PMID: 20300169] and MassHunter Quantitative Analysis B.04.00, Agilent Technologies, Santa Clara, CA, USA, respectively) in order to obtain peak information including *m/z*, migration time (MT), and peak area. The peaks were annotated with putative metabolites from the HMT metabolite database based on their MTs in CE and *m*/*z* values determined by TOFMS. The tolerance range for the peak annotation was configured at ±0.5 min for MT and ±10 ppm for *m/z*. In addition, concentrations of metabolites were calculated by normalizing the peak area of each metabolite with respect to the area of the internal standard and by using standard curves, which were obtained by three-point calibrations.

Hierarchical cluster analysis (HCA) and principal component analysis (PCA) were performed by our proprietary software, PeakStat and SampleStat, respectively.

Detected metabolites were plotted on metabolic pathway maps using VANTED (Visualization and Analysis of Networks containing Experimental Data) software [PMID:16519817].

### CE-TOFMS analysis

Total mouse T cells were isolated from spleen and lymph nodes of WT mice and were activated in 6-well plates by plate-bound anti-CD3 (2 μg/mL) and anti-CD28 (2 μg/mL) antibodies with IL-2 (5 ng/mL) in conditional media (Gln, Arg, Pro triple-free RPMI-1640 medium) containing 2 mM ^13^C_5_-Glutamine, 1.15 mM ^13^C_6_-Arginine, or 0.17 mM ^13^C_5_-Proline, respectively, for 36 hours. Activated cells (around 1.8×10^7^ cells/sample) was used for the extraction of intracellular metabolites. The cells were collected by centrifugation (300 ×*g* at 4°C for 5 min) and washed twice with 5% mannitol solution (10 mL first and then 2 mL). The cells were then treated with 800 μL of methanol and vortexed for 30 s in order to inactivate enzymes. Next, the cell extract was treated with 550 μL of Milli-Q water containing internal standards (H3304-1002, Human Metabolome Technologies, Inc., Tsuruoka, Japan) and vortexed for 30 s. The extract was obtained and centrifuged at 2,300 ×*g* and 4°C for 5 min, and then 700 μL of upper aqueous layer was centrifugally filtered through a Millipore 5-kDa cutoff filter at 9,100 ×*g* and 4°C for 180 min to remove proteins. The filtrate was centrifugally concentrated and re-suspended in 50 μL of Milli-Q water for CE-MS analysis. Metabolome measurements were carried out through a facility service at Human Metabolome Technology Inc., Tsuruoka, Japan. Hierarchical cluster analysis (HCA) and principal component analysis (PCA) were performed by our proprietary software, PeakStat and SampleStat, respectively. Detected metabolites were plotted on metabolic pathway maps using VANTED (Visualization and Analysis of Networks containing Experimental Data) software.

CE-TOFMS measurement was carried out using an Agilent CE Capillary Electrophoresis System equipped with an Agilent 6210 Time of Flight mass spectrometer, Agilent 1100 isocratic HPLC pump, Agilent G1603A CE-MS adapter kit, and Agilent G1607A CE-ESI-MS sprayer kit (Agilent Technologies, Waldbronn, Germany). The systems were controlled by Agilent G2201AA ChemStation software version B.03.01 for CE (Agilent Technologies, Waldbronn, Germany). The metabolites were analyzed by using a fused silica capillary (50 μm *i.d.* × 80 cm total length), with commercial electrophoresis buffer (Solution ID: H3301-1001 for cation analysis and H3302-1021 for anion analysis, Human Metabolome Technologies) as the electrolyte. The sample was injected at a pressure of 50 mbar for 10 s (approximately 10 nL) in cation analysis and 25 s (approximately 25 nL) in anion analysis. The spectrometer was scanned from *m/z* 50 to 1,000. Other conditions were as described previously [PMID:10740865, PMID:12038746, PMID:14582645].

### Statistical analysis

Statistical analysis was conducted using the GraphPad Prism software (GraphPad Software, Inc.). P values were calculated with two-way ANOVA for the EAE experiments. Unpaired two tail student’s t-test was used to assess differences in all other experiments. P values smaller than 0.05 were considered significant, with p-values<0.05, p-values<0.01, and p-values<0.001 indicated as *, **, and ***, respectively.

## Acknowledgments

Acknowledgments, when needed, should include the following information in the order listed below, a single paragraph, starting with the word “**Acknowledgments:**” in bold. Please use the subhead and boldface layout as shown below.).

## General

We thank John Sherman and Hayley Rodgers for critically reading and editing the manuscript.

