## Supplemental Figure 1-6 for "De novo synthesis and salvage pathway coordinately regulates polyamine homeostasis and determines T cell proliferation and function"

857 Supplementary Materials

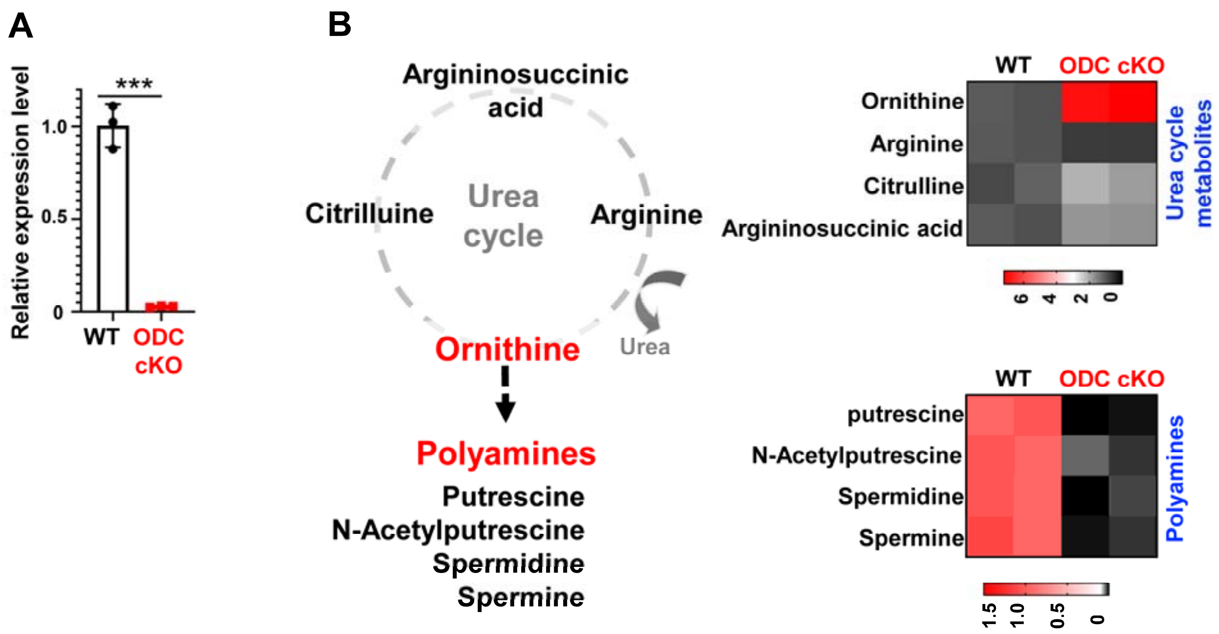


858 **Supplemental figure 1. Genetic ablation of ODC disrupts the urea cycle and polyamine pool in vitro.**

859

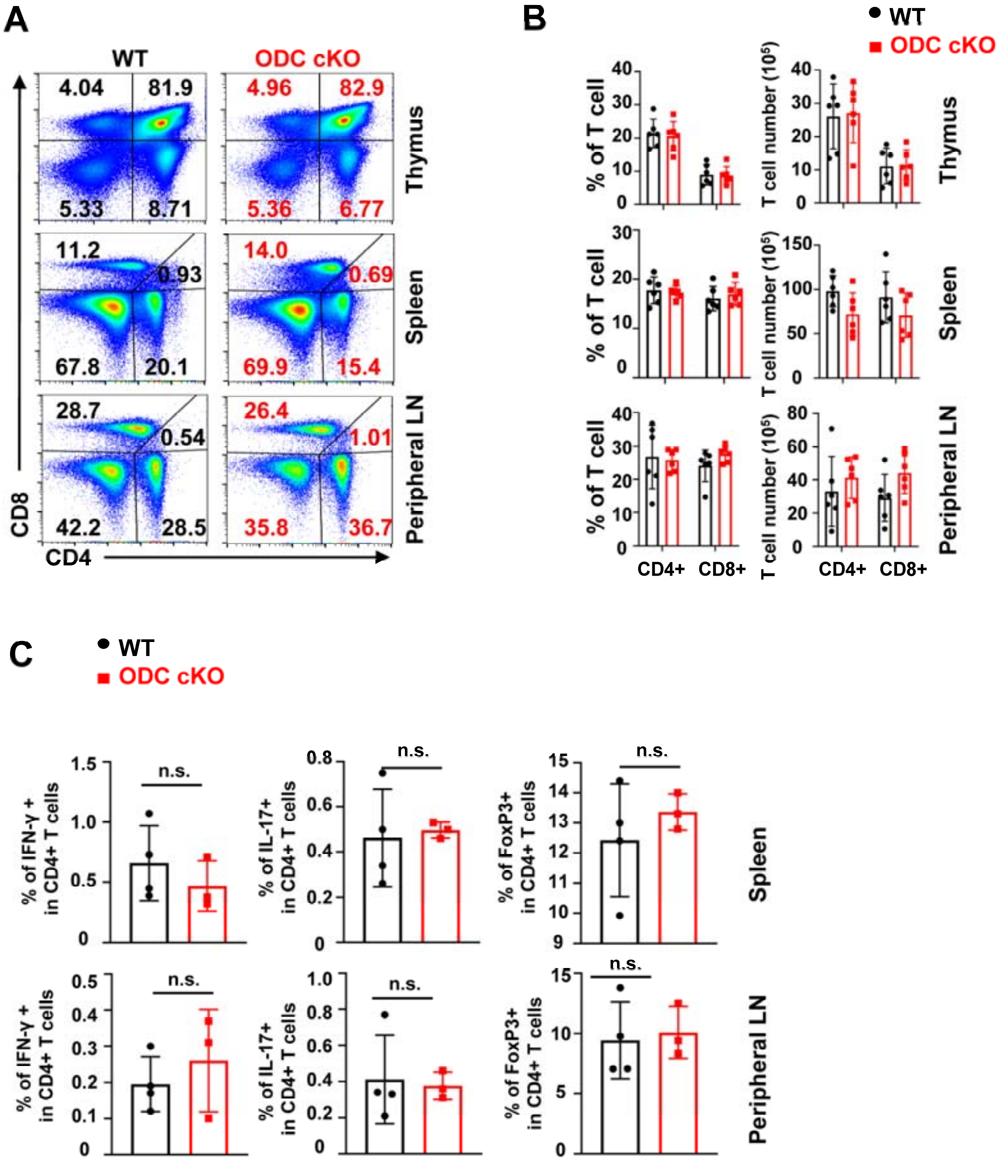


**Supplemental figure 2. ODC-mediated de novo polyamine biosynthesis is dispensable for the development of mature T cells.**

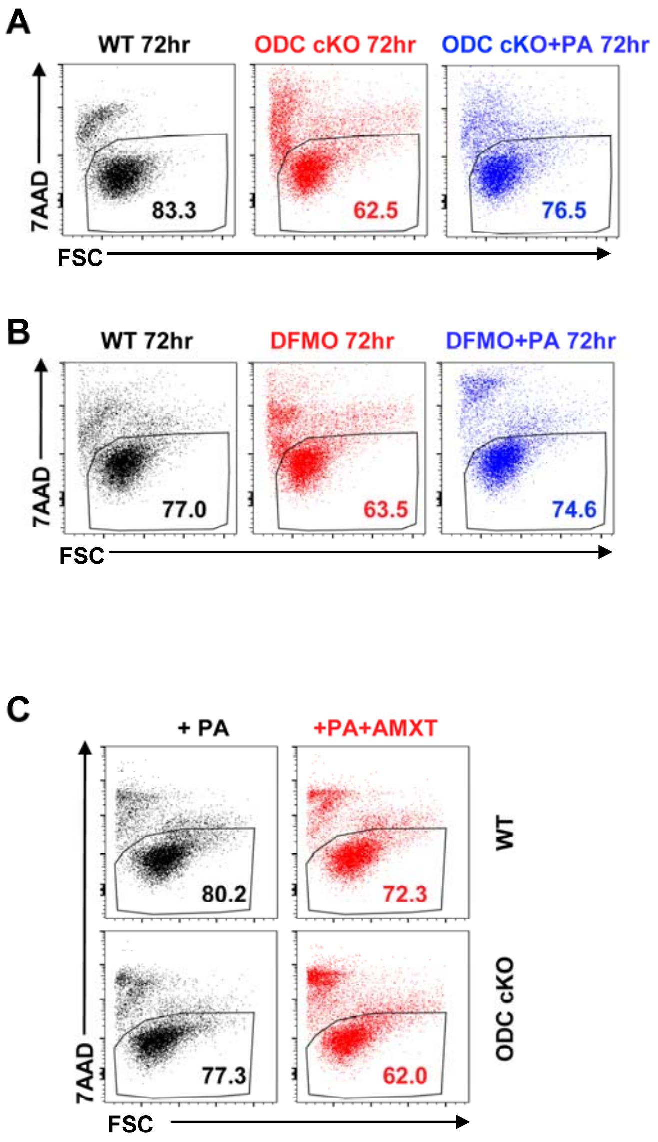


**Supplemental Figure 3. Ablation of de novo polyamine biosynthesis renders T cell dependent on polyamine uptake for survival.**

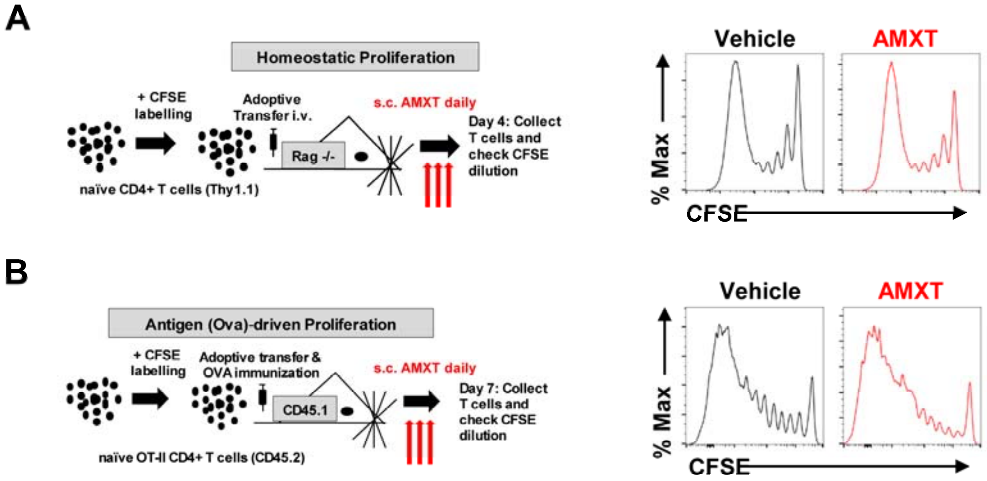


**Supplemental Figure 4. AMXT treatment doesn't block T cell proliferation in vivo.**

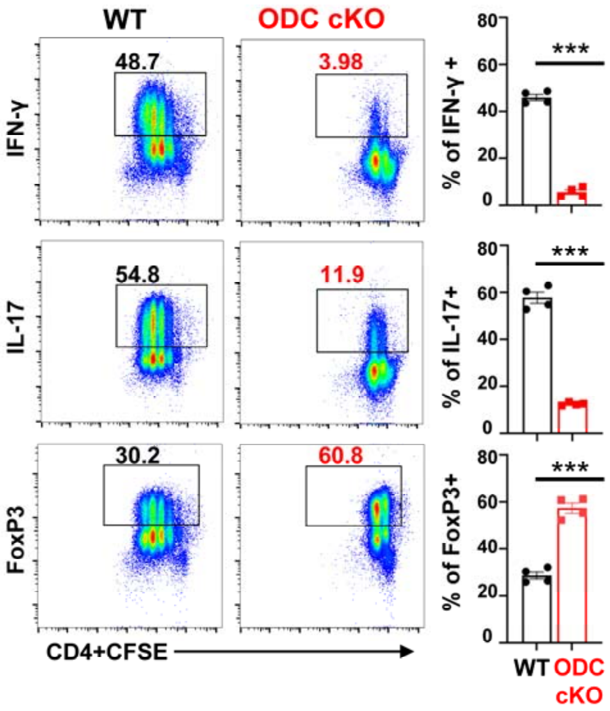


**Supplemental Figure 5. Depletion of polyamine suppresses $T_H1$, $T_H17$ but enhances $iT_{reg}$ polarization in vitro.**

867

868

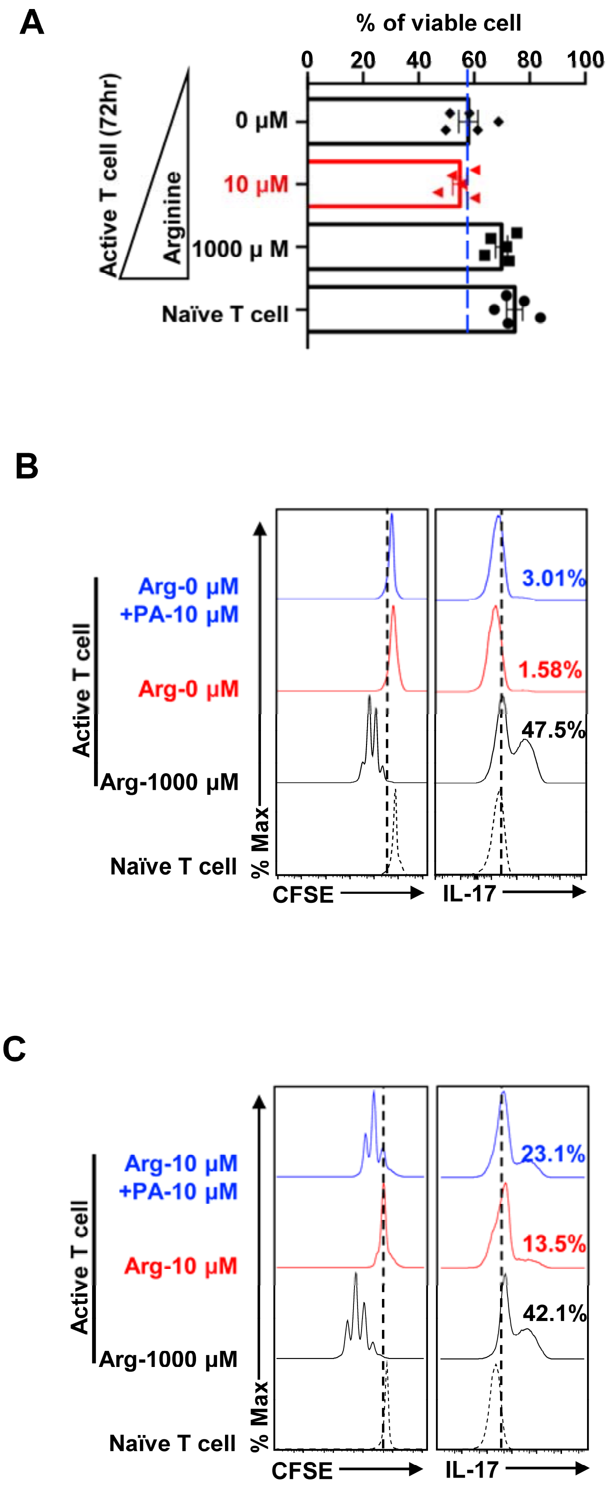


**Supplemental Figure 6. Polyamine supplement renders T cell less dependent on arginine.**

**Supplemental figure 1. Genetic ablation of ODC disrupts the urea cycle and polyamine pool *in vitro*.** (**A**), Deletion of ODC in activated ODC cKO T cells by qPCR. (**B**), schematic view of polyamine *de novo* synthesis pathway through urea cycle (left panel), and indicated intracellular metabolites in T cells collected 36 hours after plate-bound activation, profiled by **CE-QqQ/TOFMS analysis** (right panel). (**A**) representative of 3 independent experiments. (**B**) representative of 2 independent experiments. Bar graph indicates mean ± SD.

**Supplemental figure 2. ODC-mediated de novo polyamine biosynthesis is dispensable for the development of mature T cells.** (**A**), Total thymocytes, splenocytes, and lymphocytes were isolated from mice with indicated genotype. After red blood cell lysis, T cell distribution was evaluated with CD4/CD8 cell surface staining and flow cytometry. (**B**), quantification of T cell distribution and cell numbers. (**C**), endogenous $T_H1$ (IFN-ɣ expression), $T_H17$ (IL-17 expression), and $T_{reg}$ (FoxP3 expression) populations were evaluated after 4hours *ex vivo* PMA/ionomycin stimulation followed by intracellular staining and analyzed by flow cytometry. (**A-C**) representative of 3 independent experiments with each experiment containing 3-4 mice in each group. Bar graph indicates mean ± SD.

**Supplemental Figure 3. Ablation of de novo polyamine biosynthesis renders T cell dependent on polyamine uptake for survival.** (**A-C**), (A)T cells were isolated from WT or ODC cKO mice and activated using plate bound antibodies for 72 hour as indicated condition and cell viability was assessed by 7AAD uptake. (B) cell death of active T cells was evaluated in presence and absence of DFMO and DFMO supplemented with PA by 7AAD. (C) T cells were isolated from WT or ODC cKO and cultured in presence of PA mixture with and without treatment of 1 μM AMXT and cell death was evaluated by 7AAD uptake. (**A-C**) representative of 4 independent experiments.

**Supplemental Figure 4. AMXT treatment doesn't block T cell proliferation in vivo.** (**A**), CFSE-labeled donor T cells were adoptively transferred into *Rag*$^{-/-}$ lymphopenic host mice, which were randomized for treatment with vehicle (PBS) or AMXT1501 (3mg/kg/day s.c.) for 4 days. Donor cells were then retrieved, and cell proliferation was analyzed by flow cytometry. (**B**), isolated donor OTII cells labeled with CFSE were adoptively transferred into host mice, which were immunized with OVA peptides and divided into two groups for vehicle (PBS as control) or AMXT1501 (3mg/kg/day s.c.) treatment. After 7 days of *in vivo* antigen-stimulation, draining lymph nodes were isolated, and donor cell proliferation was analyzed by flow cytometry. (**A, B**) representative of 2 independent experiments.

**Supplemental Figure 5. Depletion of polyamine suppresses $T_H1$, $T_H17$ but enhances $iT_{reg}$ polarization *in vitro*.** (**A**), naïve CD4+ T cells isolated from WT and ODC cKO mice were labeled with CFSE, activated with plate-bound antibodies (anti-CD3/CD28), and differentiated in the indicated differentiation conditions for 72 hours. The indicated proteins IFN-ɣ ($T_H1$), IL-17 ($T_H17$), and FoxP3 ($iT_{reg}$) were quantified by intracellular staining following PMA and ionomycin stimulation. Representative dot plots were shown in the left panel, while the percentages of the indicated intracellular proteins in WT and ODC cKO T cells were shown in the right panel. Data are representative of 4 independent experiments. Bar graphs indicate mean $\pm$ SD.

**Supplemental Figure 6. Polyamine supplement renders T cell less dependent on arginine.** (**A-C**), T cells were isolated and maintained in culture media containing 5 ng/mL IL-7 as naïve T cells or activated in the arginine-free conditional media with 5 ng/mL IL-2 and increased concentrations of arginine (0 μM, 10 μM, and 1000 μM), respectively, with or without polyamine mixture supplement (10 μM). (**A**), cell viability was evaluated by 7AAD uptake assay in the indicated groups. representative of five independent experiments. Bar graphs indicate mean $\pm$ SD. (**B, C**) cell proliferation (CFSE) and its ability to polarize into $T_H17$ (IL-17) were evaluated at 72 hours for indicated culture condition. (**A**) Data are representative of 5 independent experiments. (**B, C**) representative of 3 independent experiments.
